# Design of T cell epitope-based vaccine candidate for SARS-CoV-2 targeting nucleocapsid and spike protein escape variants

**DOI:** 10.1101/2021.09.11.459907

**Authors:** Gabriel Jabbour, Samantha Rego, Vincent Nguyenkhoa, Sivanesan Dakshanamurthy

**Author notes:** Corresponding Author Dr. Sivanesan Dakshanamurthy, Ph.D, MBA.

## Abstract

The current COVID-19 pandemic continues to spread and devastate in the absence of effective treatments, warranting global concern and action. Despite progress in vaccine development, the rise of novel, increasingly infectious SARS-CoV-2 variants makes it clear that our response to the virus must continue to evolve along with it. The use of immunoinformatics provides an opportunity to rapidly and efficiently expand the tools at our disposal to combat the current pandemic and prepare for future outbreaks through epitope-based vaccine design. In this study, we validated and compared the currently available epitope prediction tools, and then used the best tools to predict T cell epitopes from SARS-CoV-2 spike and nucleocapsid proteins for use in an epitope-based vaccine. We combined the mouse MHC affinity predictor and clinical predictors such as HLA affinity, immunogenicity, antigenicity, allergenicity, toxicity and stability to select the highest quality CD8 and CD4 T cell epitopes for the common SARS-CoV-2 variants of concern suitable for further preclinical studies. We also identified variant-specific epitopes to more precisely target the Alpha, Beta, Gamma, Delta, Cluster 5 and US variants. We then modeled the 3D structures of our top 4 N and S epitopes to investigate the molecular interaction between peptide-MHC and peptide-MHC-TCR complexes. Following in vitro and in vivo validation, the epitopes identified by this study may be used in an epitope-based vaccine to protect across all current variants, as well as in variant-specific booster shots to target variants of concern. Immunoinformatics tools allowed us to efficiently predict epitopes in silico most likely to prove effective in vivo, providing a more streamlined process for vaccine development in the context of a rapidly evolving pandemic.

## Introduction

In late 2019, a novel coronavirus was detected in patients in Wuhan, China, and was later classified as Severe Acute Respiratory Syndrome Coronavirus 2 (SARS-CoV-2). The disease caused by SARS-CoV-2 infection, Coronavirus Disease 2019 (COVID-19), was declared a pandemic by the WHO in March 2020 and has since rapidly spread across the globe. The effects of this pandemic have been devastating, causing over 218 million infections and 4.5 million deaths worldwide as of September 2, 2021 [1], and these numbers continue to grow. While treatment options remain limited, the development of effective vaccines is crucial to curbing the spread of SARS-CoV-2 and combating the COVID-19 pandemic. The FDA has provided Emergency Use Authorizations for two vaccines as well as full approval for one vaccine, all of which are helping to lessen the toll of COVID-19 [2,3,4]. However, the rise of new, increasingly infectious variants along with a lack of global vaccine access means that the devastating effects of COVID-19 continue to be felt worldwide. It is therefore necessary to continue development of new vaccines that can combat current variants, as well as establishing a streamlined process for the rapid development of effective vaccines against novel variants. We believe that a viral epitope-based vaccine could fulfill this need as an alternative to the current vaccines. An epitope-based vaccine may be able to more precisely target certain mutations of variants of concern, as well as allowing for swift development of new vaccines through immunoinformatics as novel variants emerge in the future.

Immunoinformatics provides a more rapid, efficient, and reliable approach to vaccine design, critical to increase the speed of vaccine development in the context of a swiftly spreading pandemic. Specifically, immunoinformatics offers a cost- and time-effective strategy for designing epitope-based vaccines, which have several advantages over conventional types of vaccines. They are more specific, able to generate long-lasting immunity, and can minimize adverse reactions [5]. Given these advantages and the urgency of the current COVID-19 pandemic, we sought to predict immunogenic T cell epitopes of SARS-CoV-2 for an epitope-based vaccine candidate against the common SARS-CoV-2 variants, including the variants of concern (VOC).

T cell epitopes are recognized by Major Histocompatibility Complex (MHC) molecules on the surface of particular cells, which display the antigen to T cells, necessary for inducing an immune response. Several tools have been developed for the prediction of T cell epitopes, many of which are available on the freely available Immune Epitope Database [6]. However, the accuracy of these tools in predicting immunogenic epitopes is widely variable [7]. Therefore, to ensure that the prediction tool used in our analysis was the most accurate available on IEDB.org, we systematically analyzed the available T cell epitope prediction tools by comparing their match rate with experimentally determined immunogenic SARS-CoV-2 epitopes. This allowed us to determine the best performing epitope prediction tool for our own analysis. In addition, our analysis also validates the epitope prediction tools available on IEDB for use in predicting SARS-CoV-2 epitopes and provides guidance on how to use this resource to predict high quality SARS-CoV-2 epitopes. There are two classes of MHC molecules, MHC class I, which are expressed on all nucleated cells and activate cytotoxic T cell lymphocytes (CD8+ T cells), and MHC class II, which are expressed on antigen presenting cells and activate helper T cell lymphocytes (CD4+ T cells). To ensure our vaccine would induce both a CD8+ and CD4+ T cell response, we included epitopes that can be recognized by both MHC-I and MHC-II molecules. This is necessary to confer long-term protective immunity by inducing cytotoxic T cells and neutralizing antibodies [8]. Therefore, we used the best performing epitope prediction tools to predict epitopes of the SARS-CoV-2 Nucleocapsid (N) and Spike (S) proteins predicted to bind both MHC class I and class II.

We predicted epitopes from the SARS-CoV-2 structural proteins, namely N and S protein, for our HLA peptide-based vaccine design due to their role as major antigens of SARS-CoV-2, likely to be immunogenic. In convalescent COVID-19 patients, a robust T-cell immunity against the N protein has been detected and N protein epitopes have been found to induce responses in T cells derived from recovered patients [9]. The spike (S) glycoprotein has been reported to be a crucial surface protein of SARS-CoV-2 that facilitates viral entry into the host cell. This makes it a key target for antiviral efforts, and all current COVID-19 vaccines contain the S protein. DNA vaccines containing the S gene derived from SARS-CoV-1 have also been shown to induce T-cell responses in mice [10,11]. These proteins therefore provide ideal candidates for sources of immunogenic epitopes needed for an effective vaccine.

While the currently approved vaccines have been shown to offer effective protection against the original SARS-CoV-2 strain in clinical trials [12,13,14], multiple variants that differ from those present during clinical trials are now spreading worldwide and continue to emerge. Of current concern is the Delta (B.1.617.2) variant of SARS-CoV-2. Even countries with high vaccination rates against COVID-19, such as Israel, have seen increases in new COVID-19 cases as the variant spreads [15]. Data from Israel’s health ministry suggests that the Pfizer-BioNTech vaccine is much less effective at preventing infection by the Delta SARS-CoV-2 variant than previous strains, reporting a 39% efficacy rate compared with 95% against the original strain [15]. This raise concerns that the current vaccines, designed for previous strains, may be less effective against current and future variants. We believe it is necessary to develop vaccines that can target the mutations of variants of concern, potentially for use as variant-specific booster shots. Therefore, we sought to aid in designing these epitope-based booster vaccines that target certain variants by identifying epitopes specific to the mutations of variants of concern.

In this study, we combined careful analysis of the available epitope prediction tools, epitope prediction across multiple SARS-CoV-2 antigens, MHC binding properties, and variants, as well as HLA population coverage analysis to find immunogenic CD8 and CD4 T cell epitopes for all SARS-CoV-2 variants of concern. This epitope combination presents a candidate for use in an effective epitope-based vaccine that, following experimental validation, could aid in the fight against the COVID-19 pandemic. The variant-specific epitopes identified could also be useful in designing booster shots to target particular variants. This streamlined process may prove an efficient and reliable approach to vaccine development in fighting new outbreaks as new variants arise.

## Methods

### Retrieval of SARS-CoV-2 sequence

The SARS-CoV-2 reference N and S protein sequences (accession numbers YP_009724397 and YP_009724390 respectively) [16] were downloaded from the NCBI GenBank. The lineage-defining N and S protein mutations for the common variants: alpha variant (VOC-202012/01, B.1.1.7, UK variant), Beta variant (VOC-202012/02, B.1.351, South-African variant), Gamma variant (VOC 202101/02, P.1, Brazil-Japan variant) [17], US variants (N [18] and S [19] protein mutations), Delta variant (VUI B.1.617, Indian variant) [20] and Cluster 5 mink variant [21] were used to create the variant specific N and S lineage-defining protein sequences.

### Retrieval of N protein SARS-CoV-2 experimentally determined epitopes

#### CD8 experimental epitopes

An experimental epitope is selected if its immunogenicity has been experimentally determined and its length is between 8 and 14 amino acids [22]. We selected 67 epitopes from various papers [23-31], 6 from RCSB.org (7KGT, 7LG2, 7LG3, 7LFZ, 7KGR and 7KGS) and 15 epitopes from IEDB.org and ViPR.org with positive T cell assay and 100% match with the SARS-CoV-2 N reference sequence, for a total of 88 high quality epitopes with experimentally determined immunogenicity (Supplementary Material 1).

#### CD4 experimental epitopes

To find experimentally determined epitopes, we searched for SARS-CoV-2 N protein epitopes whose immunogenicity had been validated experimentally. We selected 28 epitopes from previous papers that had been found to be immunogenic in convalescent COVID-19 patients [27, 31-33] and 20 epitopes from ViPR with positive assay and 100% match with the SARS-CoV-2 N protein reference sequence. We also searched the RCSB Protein Data Bank for known SARS-CoV-2 N protein epitopes binding to MHC class II, which returned no results. This provided a total of 48 high quality epitopes with experimentally determined immunogenicity (Supplementary Material 1).

### Choosing the best tool

The 5 common CD8 T cell epitope prediction tools available on IEDB: ANN 4.0 [34-39], IEDB consensus [40], IEDB recommended for proteasome-TAP-MHC1 binding combined predictor (IEDB Prot-TAP-MHC) [41,42], NetMHCpan EL 4.1 (IEDB recommended 2020.09) and NetMHCpan BA 4.1 [43-47] and the 4 common CD4 T cell epitope prediction tools available on IEDB: IEDB recommended 2.22, IEDB consensus 2.22 [48,49], NetMHCIIpan EL 4.0 and NetMHCIIpan BA 4.0 [50,51] were used to predict SARS-CoV-2 N protein epitopes. Only MHC alleles with a frequency > 1% were selected. For MHCII epitope prediction, the default size of 15 amino acids per epitope was selected.

The duplicate-free peptides were first ordered according to each of the available attributes in each tool, then matched to the experimental epitopes according to 3 stringency levels: lenient, intermediate and stringent (Supplementary Material 1). Receiver operating characteristic (ROC) analysis was used to assess the quality of select peptide prediction tools. ROC evaluates binary data as positive and negative values, which was achieved by having defined thresholds based on the predicted peptides’ binding affinity. It is important to note that the measure of binding affinity (rank, IC50, score, or percentile rank) varied amongst prediction tools, which was an additional consideration when determining the best tool. The positive and negative experiment test set was obtained by assessing each tool based on the matches of predicted epitopes to experimentally determined epitopes. More specifically, a match between a predicted and experimental epitope based on the Lenient, Intermediate, or Stringent method was a positive (1), while a non-match was a negative (0).

To determine the quality of each prediction tool, specificity and sensitivity calculations were needed for the construction of the ROC curves. The tool that maximizes both specificity and sensitivity would be regarded as the best tool. This is more easily quantified by measuring the area under the curve (AUC), where an area of 0.5 indicates an entirely random prediction and an area of 1.0 indicates a perfect prediction. All analysis and data visualizations were constructed from R scripts. The pROC package was used for ROC and AUC calculations.

### Epitope prediction

Using the best tool-attribute combination, we predicted CD8 and CD4 T cell epitopes for both N and S proteins for each variant.

### MHCI epitope immunogenicity prediction

Immunogenicity scores for the top 300 epitopes from each variant were calculated using the MHC I immunogenicity tool in IEDB.org [52]. Only CD8 common epitopes with an immunogenicity score > 0 and CD8 variant-specific epitopes with an immunogenicity score > - 0.5 were selected for further analysis.

### MHCII epitope antigenicity prediction

Vaxijen 2.0 [53-55] was used to select the best CD4 epitopes due to the lack of a reliable immunogenicity prediction tool on IEDB.org.

### Toxicity prediction

Using ToxinPred [56], CD8 and CD4 peptides were found to be either toxic or non-toxic. Only non-toxic epitopes were selected for further studies.

### Allergenicity prediction

AllerTop 2.0 [57,58] was used to predict allergenicity of both CD8 and CD4 epitopes. Only “probable non-allergenic” epitopes were selected.

### Instability index prediction

ProtParam [59] was used to predict the instability index [60] to determine whether an epitope was stable or unstable in vivo.

### Mouse MHC affinity prediction

NetH2pan 4.0 [61,62] was used to predict mouse MHC affinity of CD8 epitopes using MHCI alleles of the most commonly used mouse strains to study COVID-19 vaccines: C57BL/6 and BALB/c rodents. Only Weak Binding and Strong Binding epitopes were selected for potential further preclinical studies.

### Population coverage analysis

The population coverage analysis tool available on IEDB.org [63] was used to predict the percentage of the world population predicted to present the epitopes with known MHC restrictions.

### p-MHC 3D structure prediction

The tertiary structure of the selected epitopes was predicted using PEPstrMOD [64,65]. The 3D structures of MHC molecules were downloaded from Protein Database Bank on RCSB.org and processed on PyMOL to remove the epitope. The predicted epitope along with the empty MHC molecule were energy minimized using Swiss-PdbViewer [66]. The two molecules were then docked using FlexPepDock online server [67,68].

## Results

### Choosing the best tool

For the prediction of common SARS-CoV-2 CD8 epitopes for both N and S proteins, the results suggested the use of IEDB recommended (NetMHCpan EL 4.1) ordered according to “rank”, as opposed to “score” as recommended by IEDB. However, this difference was not statistically significant. IEDB recommended ranked first in AUC at 0.7423 and 0.9192 for the Lenient and Stringent method, respectively (Figure 2). For the prediction of common SARS-CoV-2 CD4 epitopes, there was no statistically significant difference between the tools, so we decided to use the IEDB recommended ordered according to “rank,” since it was a top performer with AUC of 0.6969, 0.7078 and 0.767 for the Lenient, Intermediate and Stringent methods respectively (Figure 3).

**Figure 1.**
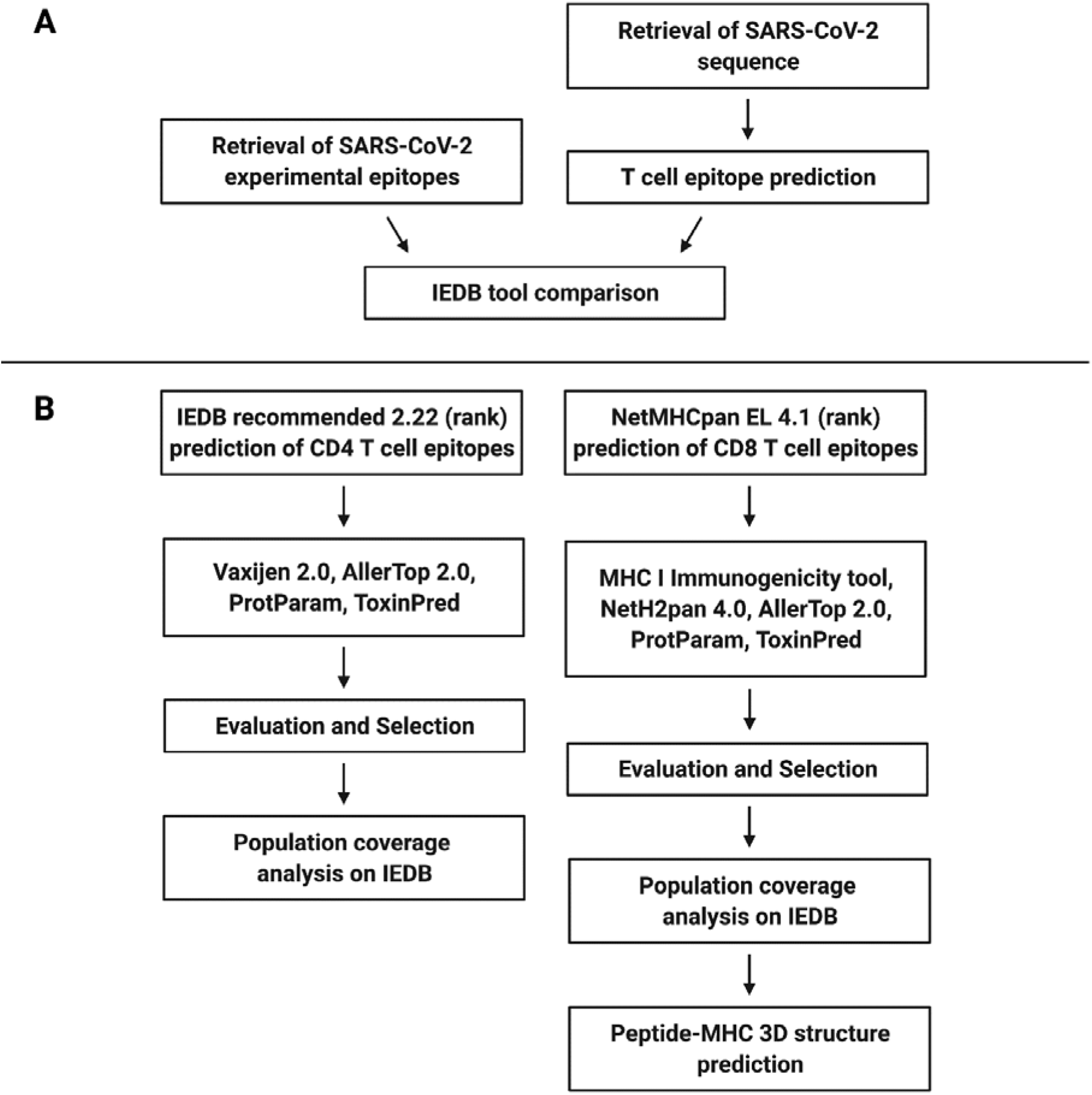
Schematic flow chart diagram of the in silico peptide prediction process. (A) Selection of the best MHC-I and -II binding epitope prediction tools based on its classification performance. (B) In silico identification and evaluation of novel CD4 and CD8 peptides for vaccine design.

**Figure 2.**
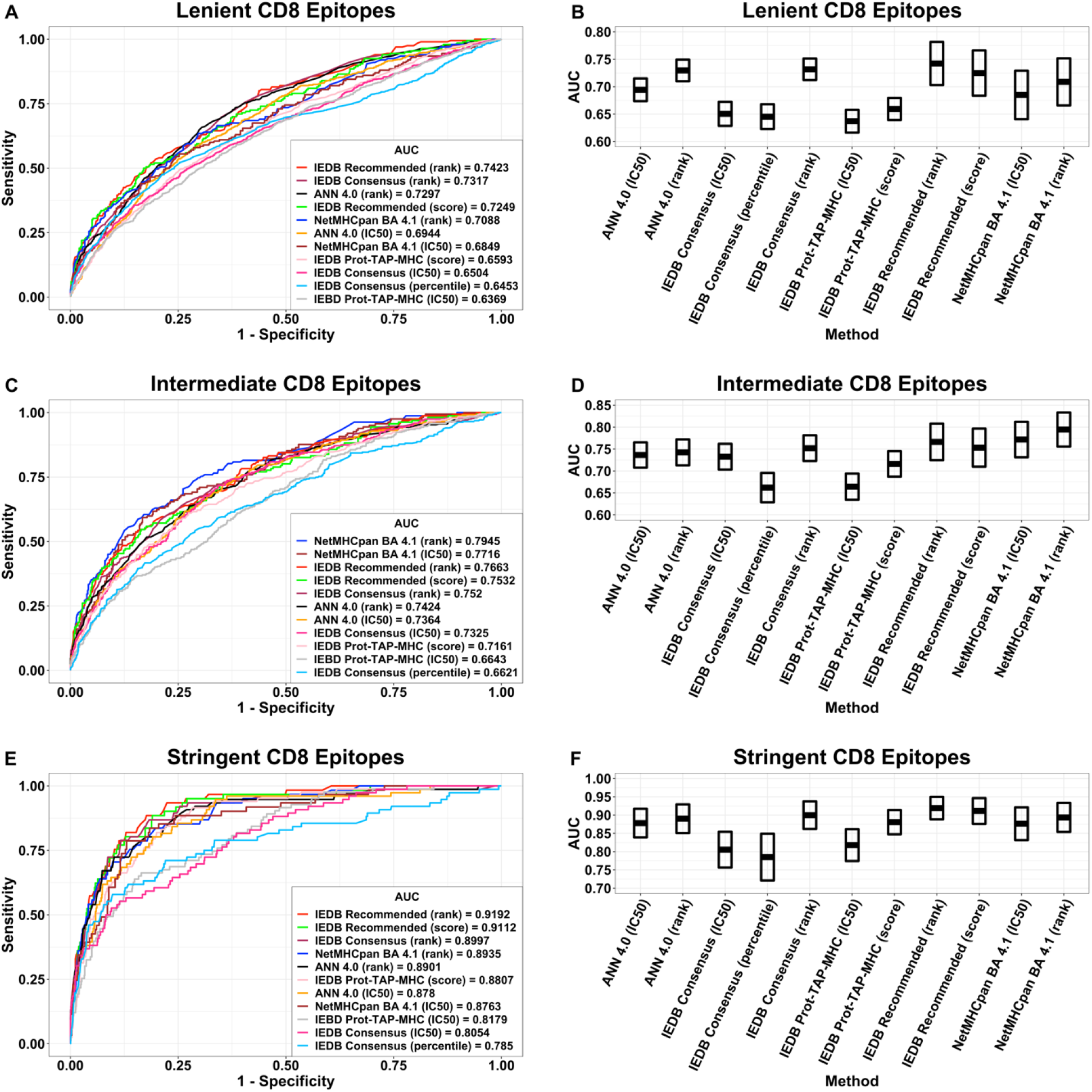
Comparison of MHC class I binding epitope prediction tools’ binary classification performance. (A, C, E) ROC curves with AUC values were shown for each tool-attribute combination that was evaluated for its experimental epitope prediction power. Cutoff thresholds for classification were established through the attribute, ordered from highest to lowest binding affinity, for each tool. (B, D, F) Boxplots show the upper and lower 95% confidence interval boundaries of each tool-attribute’s AUC values.

**Figure 3.**
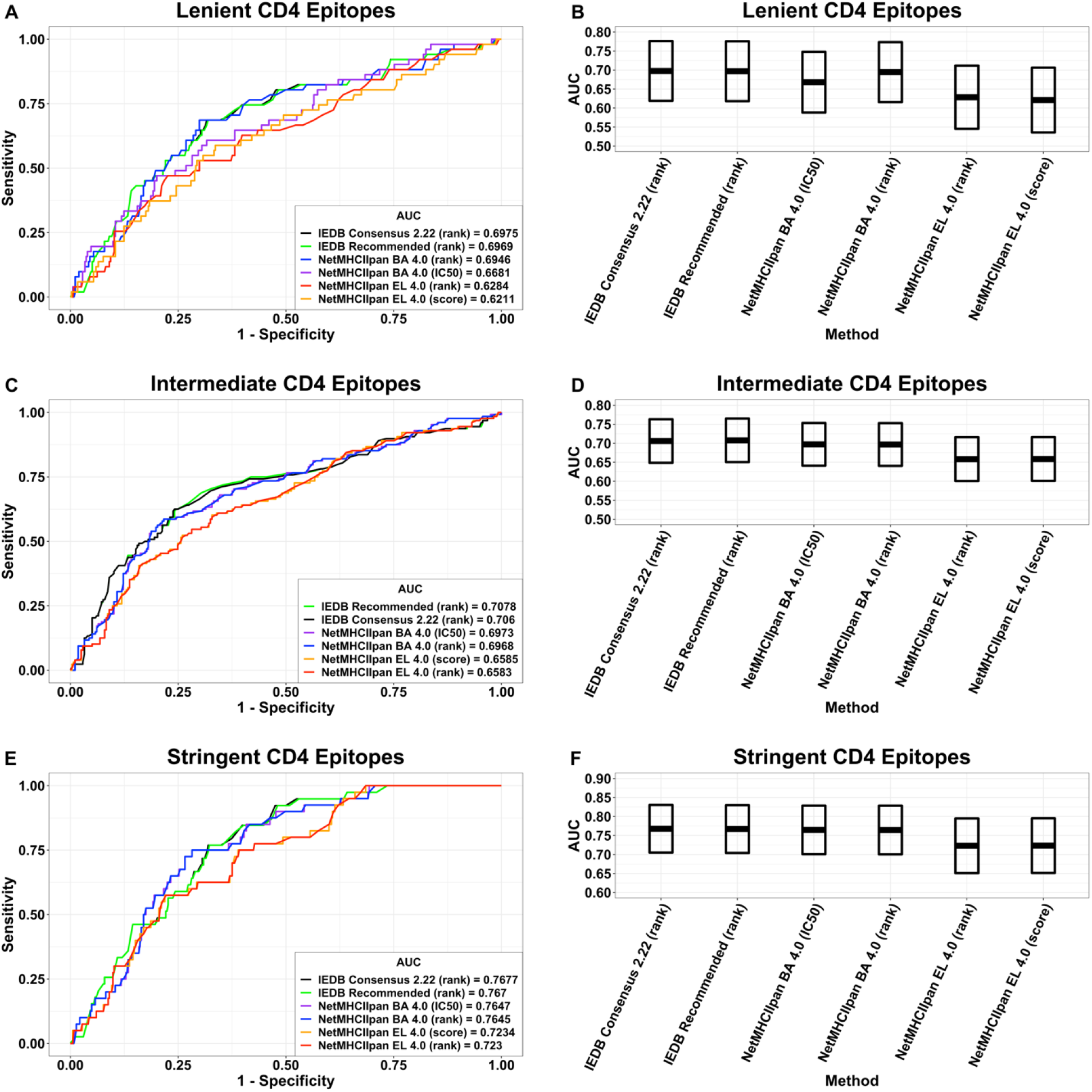
Comparison of MHC class II binding epitope prediction tools’ binary classification performance. (A, C, E) ROC curves with AUC values were shown for each tool-attribute combination that was evaluated for its experimental epitope prediction power. Cutoff thresholds for classification were established through the attribute, ordered from highest to lowest binding affinity, for each tool. (B, D, F) Boxplots show the upper and lower 95% confidence interval boundaries of each tool-attribute’s AUC values.

### CD8 epitope prediction

Using NetMHCpan EL 4.1, MHCI immunogenicity predictor, ToxinPred, AllerTop 2.0, we only selected CD8 epitopes that were immunogenic, non-toxic and non-allergenic. We obtained 31 common N and 31 common S protein epitopes across all variants (Supplementary Table S1), 1 N and 2 S epitopes specific for the beta variant, 2 N and 4 S epitopes specific for the gamma variant, 2 N and 4 S epitopes specific for the alpha variant, 3 N epitopes specific for the US-Australian variants, 2 S epitopes specific for the US variant, 2 S epitopes specific for the Cluster 5 variant and 5 S epitopes specific for the delta variant and 1 S epitope specific for both the delta and US variants (Supplementary Table S3 and Supplementary Material 2).

Adding the stability filter, we obtained 15 N and 21 S CD8 epitopes in total (Supplementary Table S2), 1 N epitope specific for the alpha variant, 1 S epitope specific for the beta variant, 3 S epitopes specific for the gamma variant, 4 S epitopes specific for the alpha variant, 2 S epitopes specific for the US variant, 2 S epitopes specific for the Cluster 5 variant, 1 S epitope specific for the delta variant and 1 S epitope specific for both the US and delta variants (Table 2).

### CD4 epitope prediction

Using IEDB recommended 2.22, Vaxijen 2.0, ToxinPred and AllerTop 2.0, we only selected the antigenic non-toxic and non-allergenic CD4 epitopes. We obtained 5 common N and 11 common S protein epitopes across all variants (Supplementary Table S4), 1 S epitope specific for the beta variant, 3 N and 8 S epitopes specific for the gamma variant, 4 S epitopes specific for the alpha variant, 1 N epitopes specific for the US-Australia variant, 5 S epitopes specific for the US variant, 3 N and 7 S epitopes specific for the delta variant, 1 S epitope common to the delta and US variants, and 3 S epitopes specific for the Cluster 5 variant (Supplementary Table S5 and Supplementary Material 3).

With stability prediction, we obtained 1 common N and 7 common S epitopes (Table 3), 1 S epitope specific for the beta variant, 7 S epitopes specific for the gamma variant, 1 S epitope specific for the alpha variant, 4 S epitopes specific for the US variant, 3 N and 3 S epitopes specific for the delta variant, 1 S epitope common to the delta and US variants, and 3 S epitopes specific for the Cluster 5 variant (Table 4).

### Population coverage analysis

The 31 N and 31 S CD8 epitopes cover 98.46% and 92.02% of the world population respectively. However, since epitope-based vaccine constructs usually contain fewer epitopes, we also found the regional and world coverage of our top 11 N and top 15 S CD8 epitopes to be 97.87% and 89.92% respectively (Table 1, Figure 4 and Supplementary Material 4). These 23 highly immunogenic, non-toxic and non-allergenic peptides are predicted to elicit a protective immune response in 98.42% of the world population. After adding the instability index prediction and filtering out the unstable epitopes, the population coverage decreased to 78.28% and 87.13% for N and S protein epitopes respectively (Supplementary Table S2).

**Table 1.**
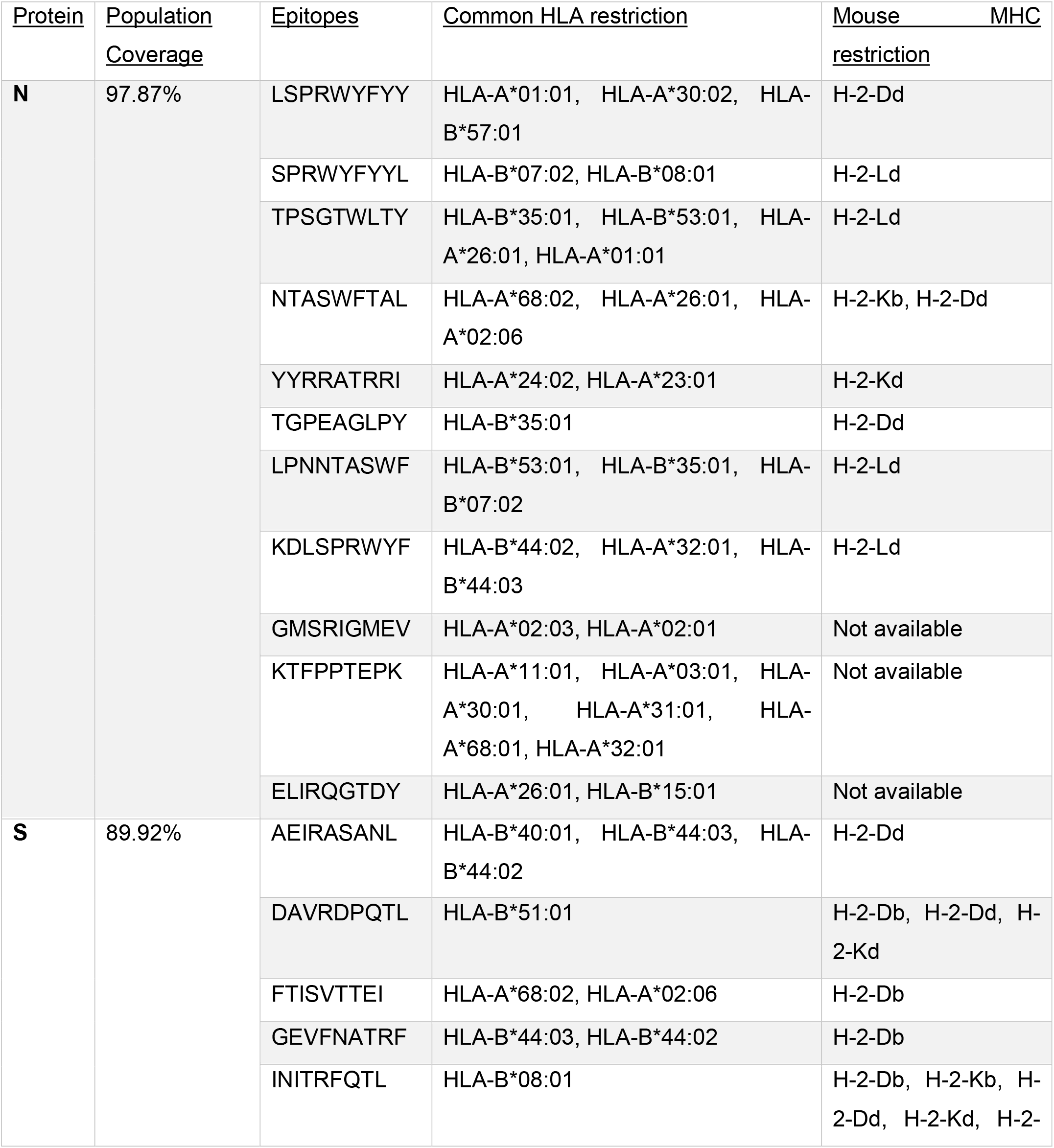
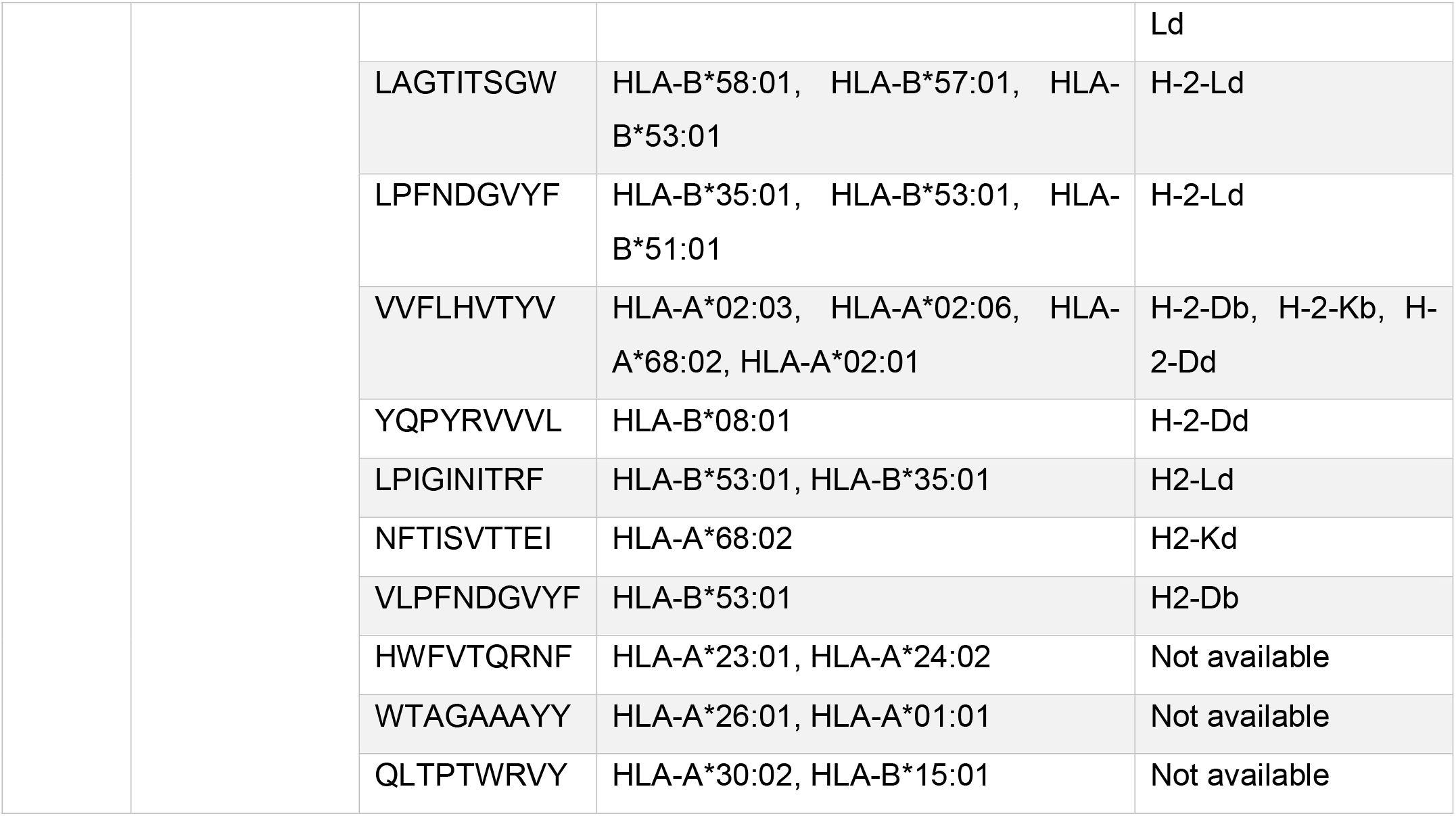
Top 11 N and top 15 S protein CD8 epitopes that are most immunogenic, non-toxic, non-allergenic and common across all studied SARS-CoV-2 variants with their world population coverage and their mouse MHC haplotype when available.

**Figure 4.**
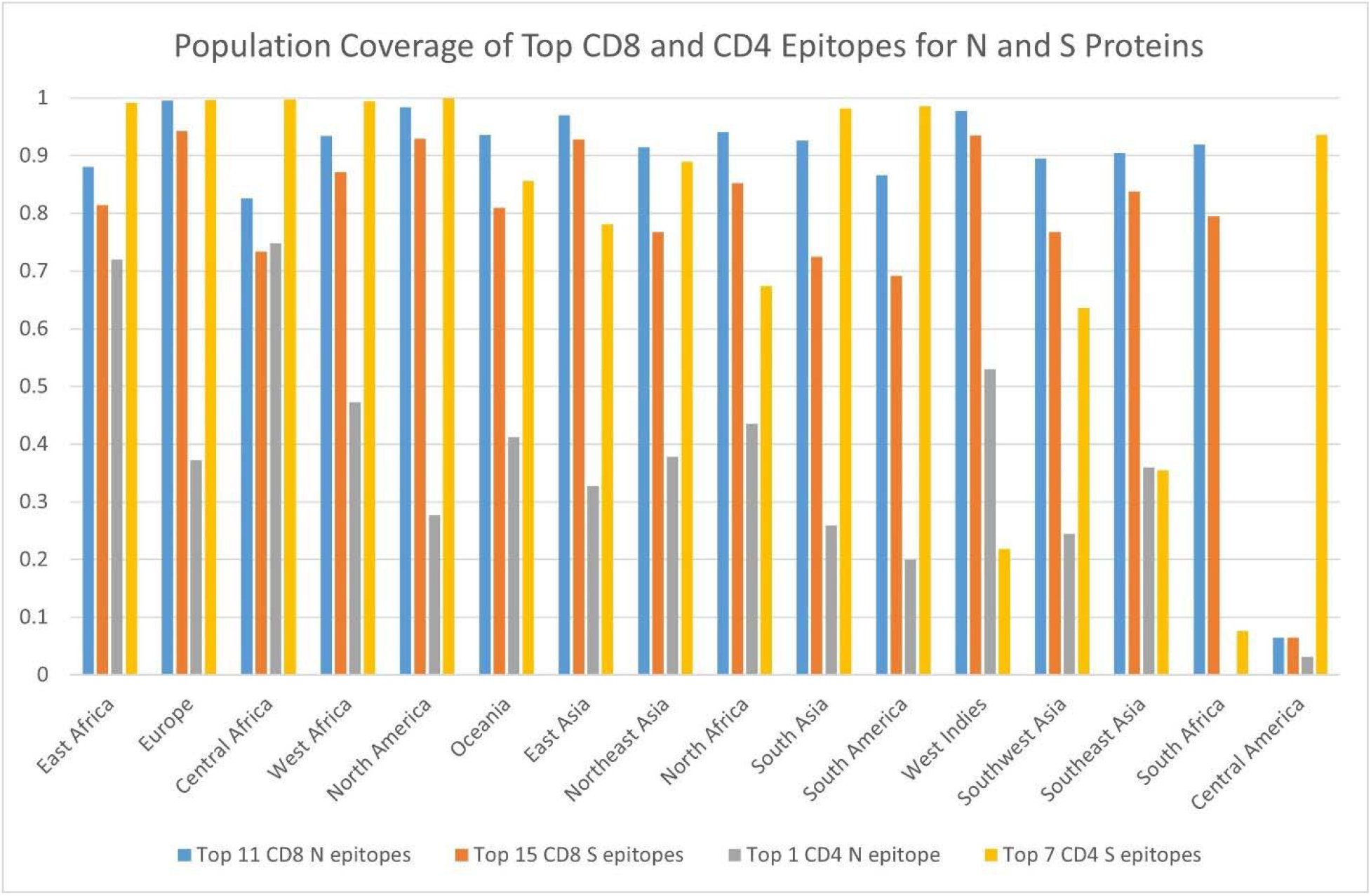
Regional population coverage of the top 11 N CD8 epitopes, top 15 S CD8 epitopes, top 1 N CD4 epitope and top 7 S CD4 epitopes.

The 5 N and 11 S CD4 epitopes have world population coverages of 45.40% and 97.11% respectively (Supplementary Table S4). After adding the stability prediction, the top 1 N and top 7 S CD4 epitopes cover 34.55% and 95.13% of the world population respectively (Table 3 and Figure 4). Together, the top 8 N and S CD4 epitopes cover 96.81% of the world.

### p-MHC 3D structure prediction

Next, 3 N (NTASWFTAL, TPSGTWLTY and LPNNTASWF) and 1 S (VVFLHVTYV) immunogenic, non-toxic, non-allergenic common CD8 epitopes with significant affinity to at least one mouse MHC haplotype were selected for 3D modeling. These epitopes were docked with their respective predicted HLA alleles: NTASWFTAL-HLA-A^*^68, TPSGTWLTY-HLA-A^*^01:01, LPNNTASWF-HLA-B^*^07:02 and VVFLHVTYV-HLA-A^*^02:01 (Figure 5). Furthermore, the VVFLHVTYV-HLA-A^*^02:01 complex was superimposed with the 7N1E RCSB structure to get an insight into the interaction between the p-MHC complex and the pRLQ3 TCR molecule (Figure 5 and Supplementary Figure S1). To avoid clash the p-MHC-pRLQ3 TCR complex was visually analyzed, and energy-minimized with the SANDER module of AMBER [69].

**Figure 5.**
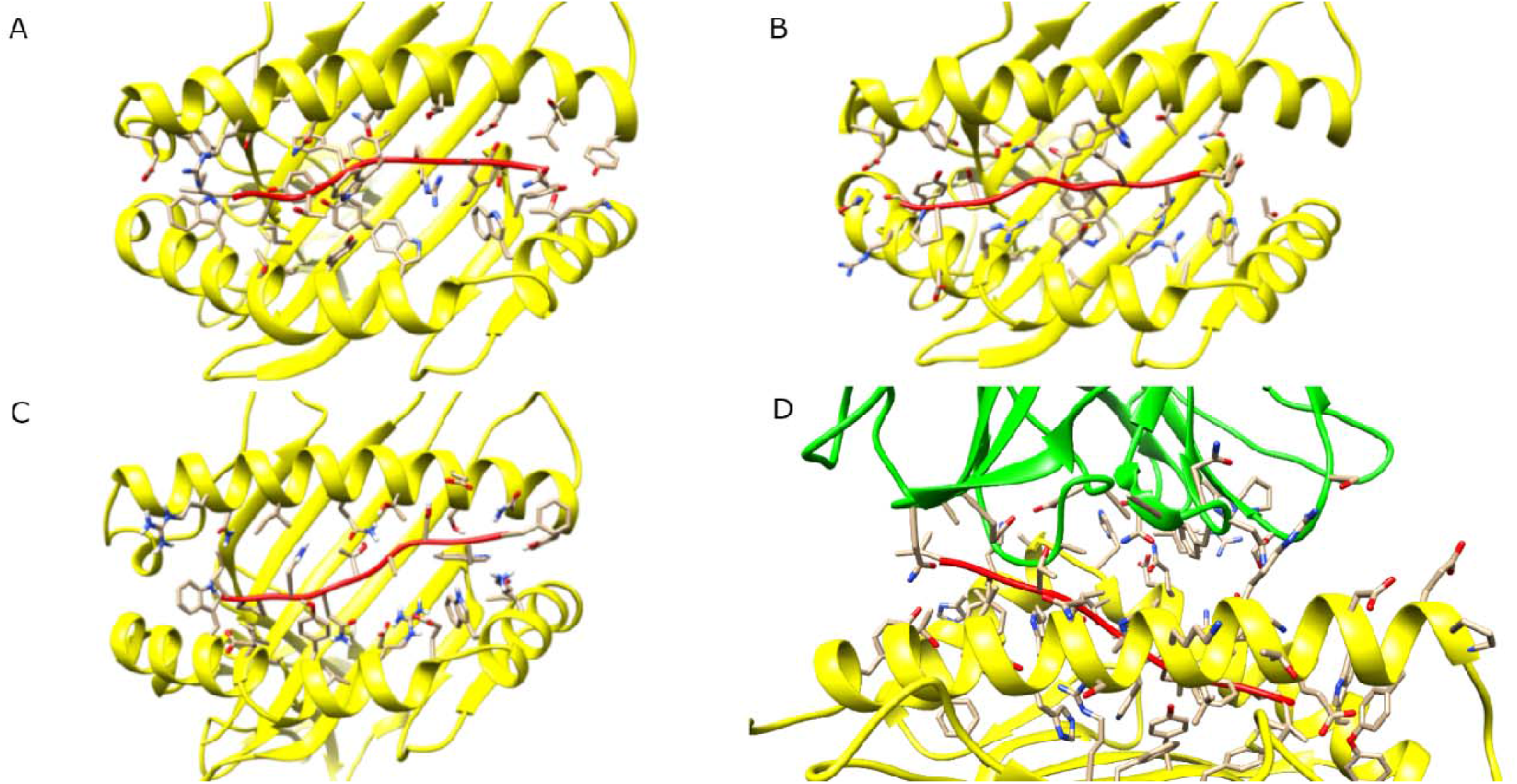
Close-up shots of the predicted 3D structure for the interaction of 3 SARS-CoV-2 N epitopes (red) with their selected MHC I molecules (yellow): NTASWFTAL-HLA-A^*^68 **(A)**, TPSGTWLTY-HLA-A^*^01:01 **(B)**, LPNNTASWF-HLA-B^*^07:02 **(C)** and 1 SARS-CoV-2 S epitope (red) with its selected HC (yellow) and TCR molecule (green): VVFLHVTYV-HLA-A^*^02:01-pRLQ3 TCR **(D)**.

## Discussion

Immunoinformatics provides an efficient and reliable strategy for designing epitope-based vaccines to combat the evolving COVID-19 pandemic, especially as new SARS-CoV-2 variants emerge and spread. However, with the ever-increasing number of machine learning tools in the field of immunoinformatics, the selection of epitope prediction tools is becoming increasingly arbitrary. It is necessary to validate the performance of a given tool before its predictions can be utilized with confidence. Our analysis of the performance of multiple epitope prediction tools against experimentally validated SARS-CoV-2 epitopes allowed us to identify the best IEDB tool for prediction of SARS-CoV-2 epitopes. We showed that among the commonly used IEDB prediction tools, NetMHCpan EL 4.1 and CD4 IEDB recommended 2.22 are ordered according to “rank”, are the most reliable combinations to predict SARS-CoV-2 CD8 and CD4 epitopes, respectively. More studies are needed to calculate a rank threshold for the epitope prediction tools to prevent the arbitrary selection of thresholds for high affinity epitope selection.

Next, we used immunoinformatics tools to predict immunogenic T cell epitopes of SARS-CoV-2 for use in a multi-epitope vaccine against the common SARS-CoV-2 variants, including the variants of concern (VOC). Our study identified 31 common N and 31 common S CD8 epitopes (Supplementary Table S1), and 5 common N and 11 common S CD4 epitopes (Supplementary Table S4), all of which are predicted to be immunogenic (or antigenic for CD4 epitopes), non-toxic and non-allergenic. Epitopes showed a world population coverage of 97.87% for the selected top 11 N, 89.92% for the selected top 15 S CD8 epitopes (Table 1), 34.55% for the selected top 1 N CD4 epitope and 95.13%% for the selected top 7 S CD4 epitopes (Table 3). These epitopes present ideal candidates for use in a candidate vaccine that could provide protection across current SARS-CoV-2 variants. B cell epitopes will also need to be identified to complete the vaccine formulation. Additionally, we identified variant-specific immunogenic, non-toxic, non-allergenic, and stable CD8 (Table 2) and CD4 (Table 4) epitopes that can more pointedly target the mutations particular to each variant of concern. Given the rise of increasingly infectious variants, as well as concerns of decreasing vaccine efficacy against new variants, these epitopes may be useful in developing variant-specific booster shots to provide stronger protection against variants.

**Table 2.**
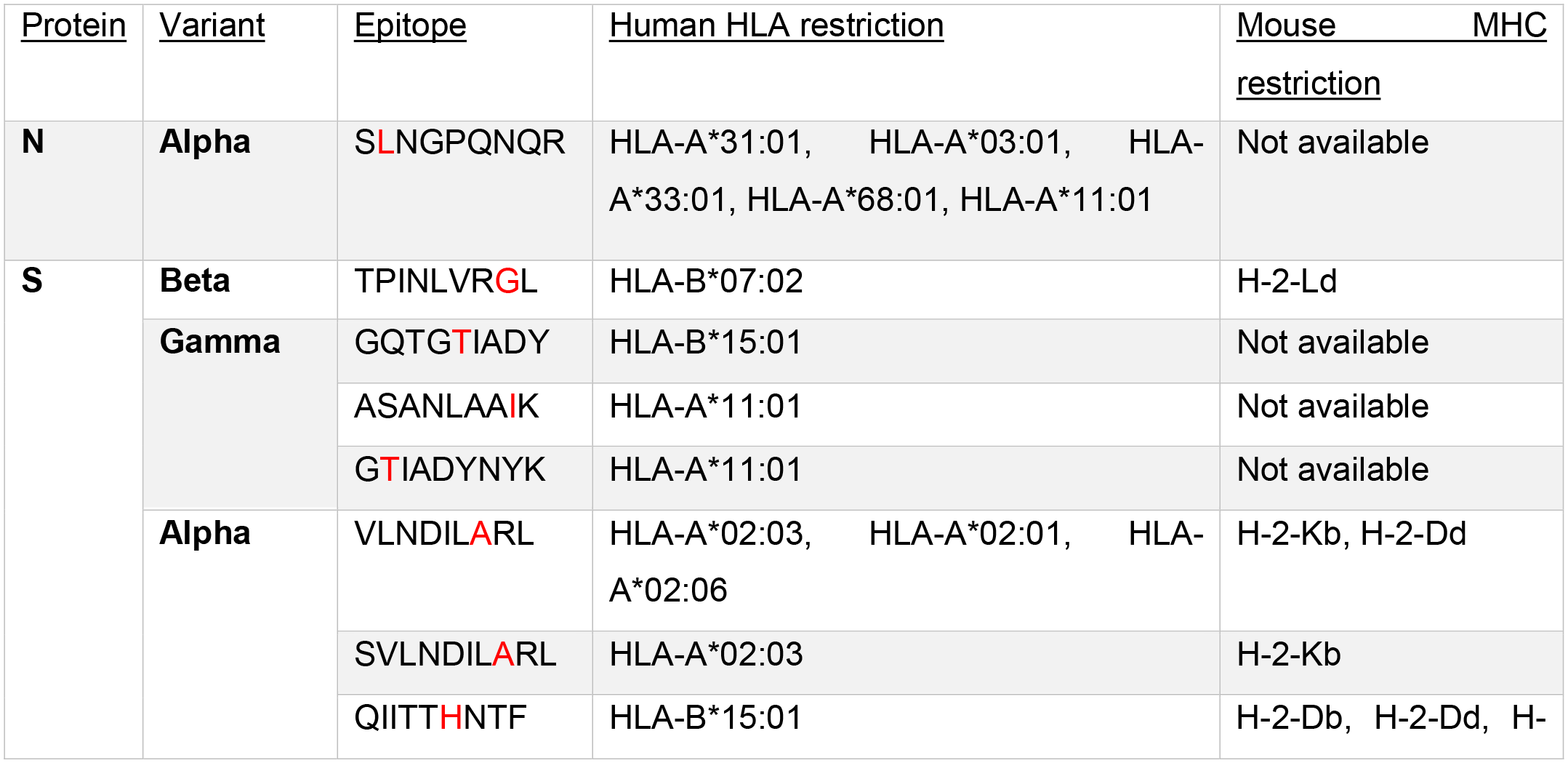
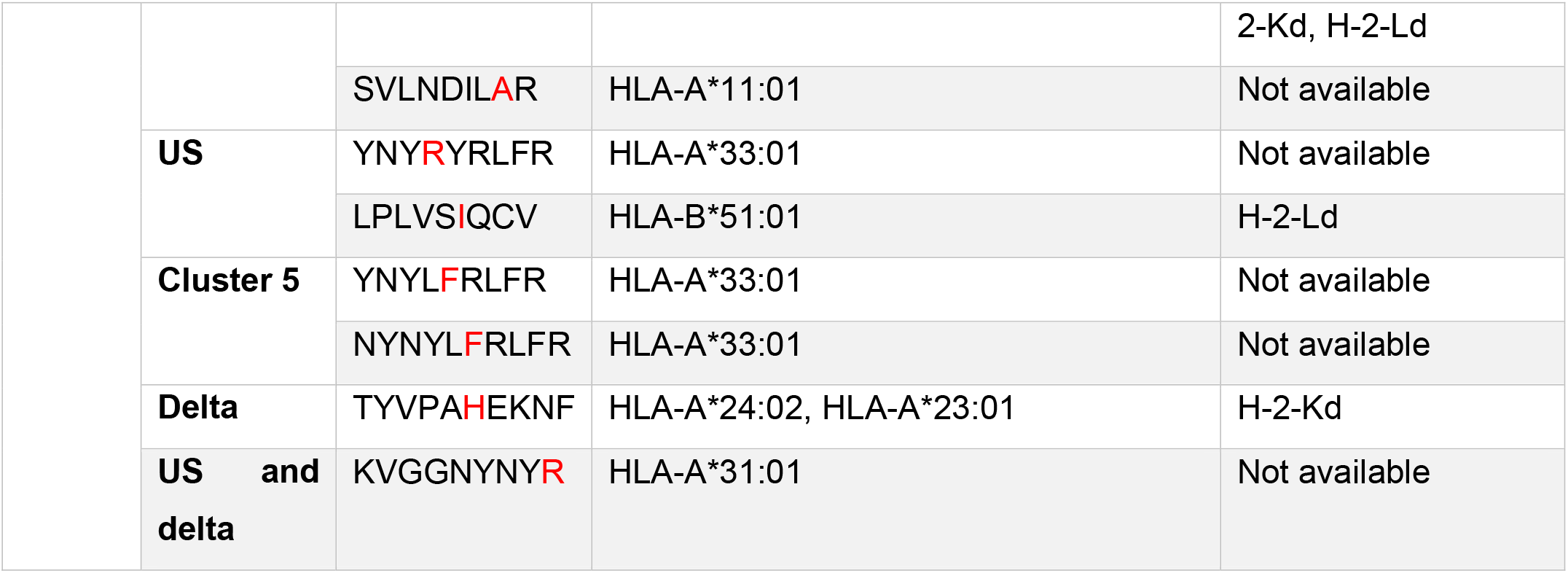
All immunogenic non-toxic non-allergenic stable CD8 variant-specific epitopes for each SARS-CoV-2 variant N and S protein, with the variant-specific mutations written in red (see Supplementary Table S3 for full list of immunogenic non-toxic non-allergenic variant-specific CD8 epitopes).

**Table 3.**
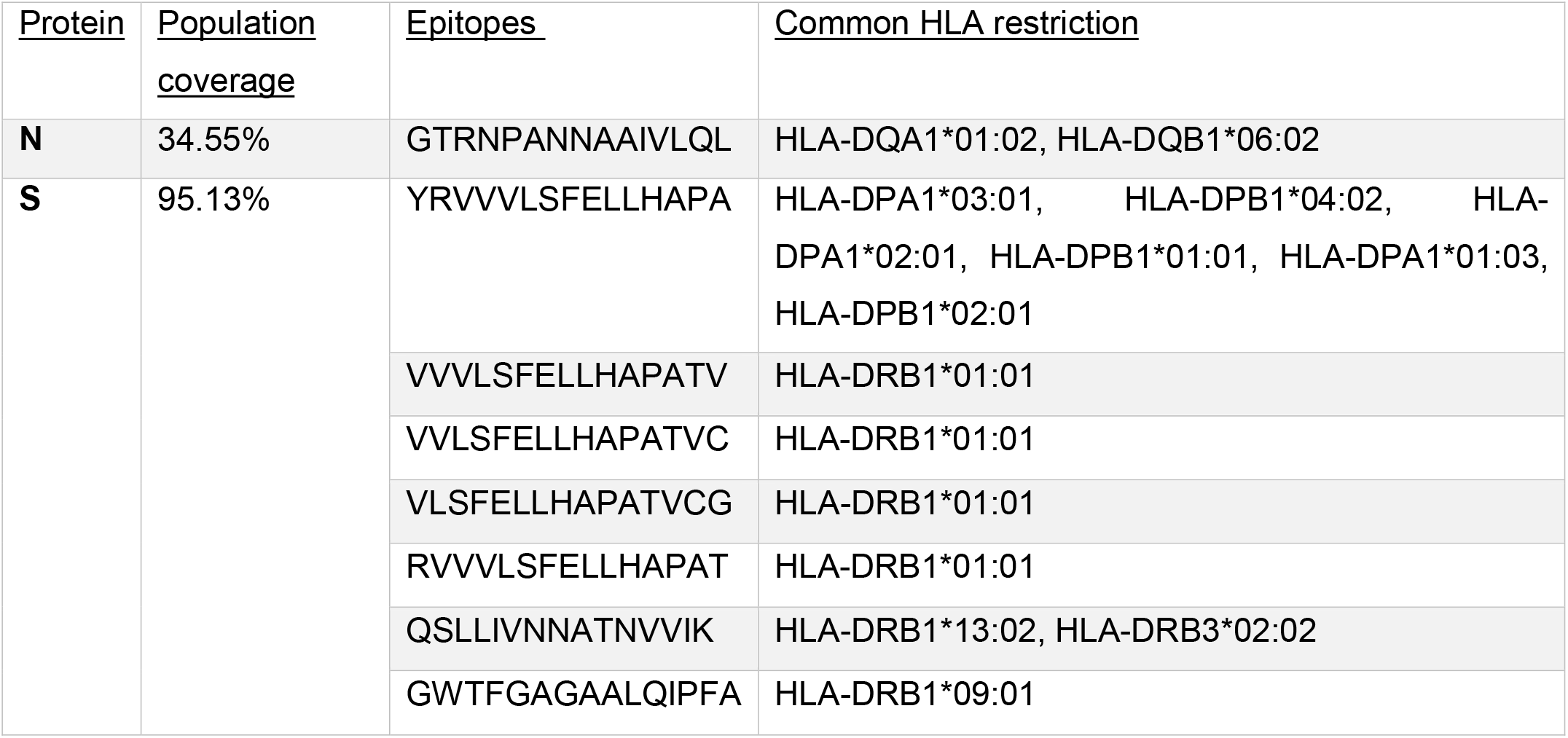
All common N and S CD4 antigenic non-toxic non-allergenic stable epitopes across all studied SARS-CoV-2 variants and their world population coverage.

**Table 4.**
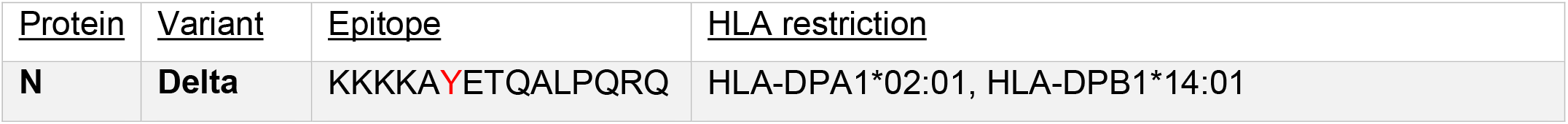
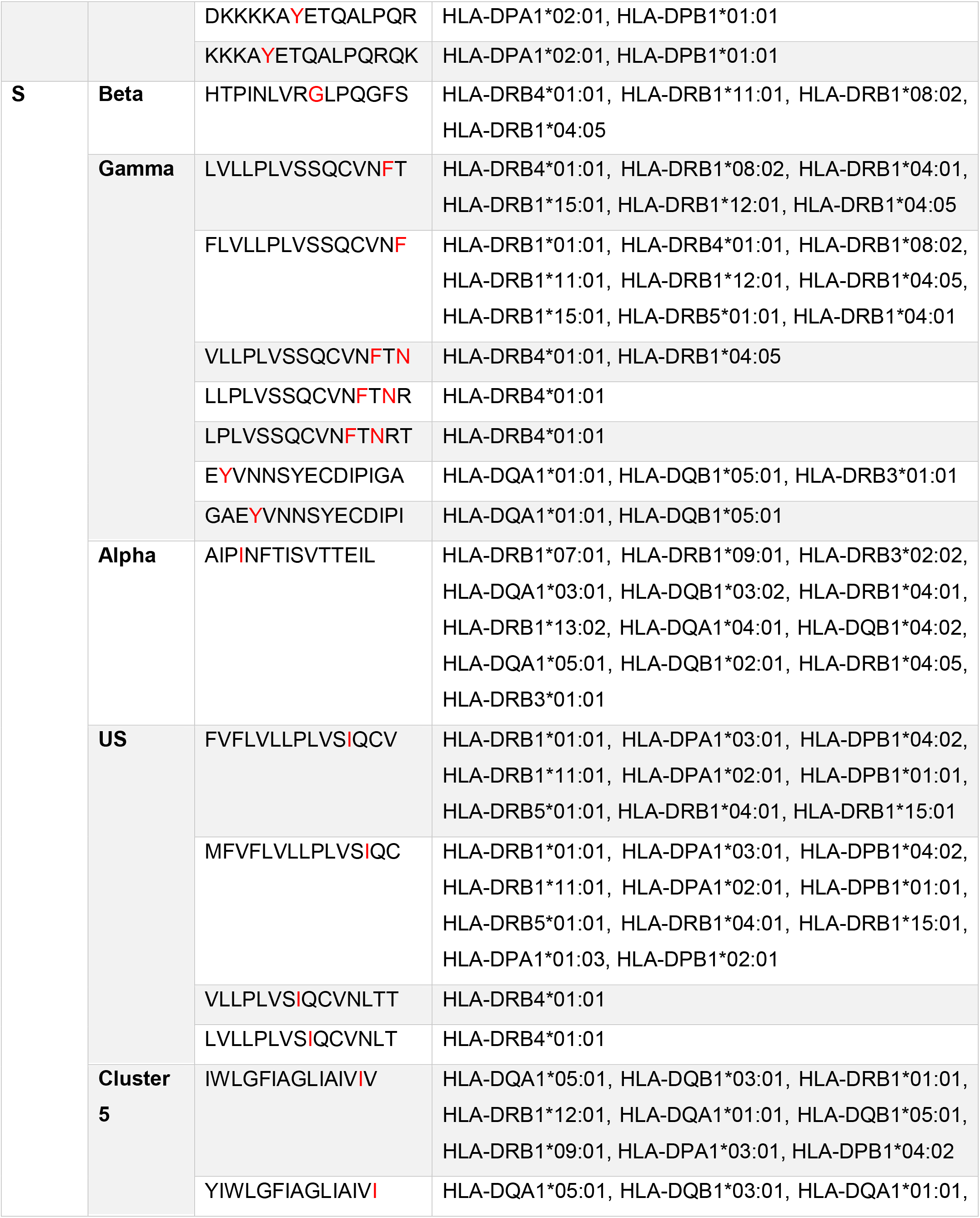
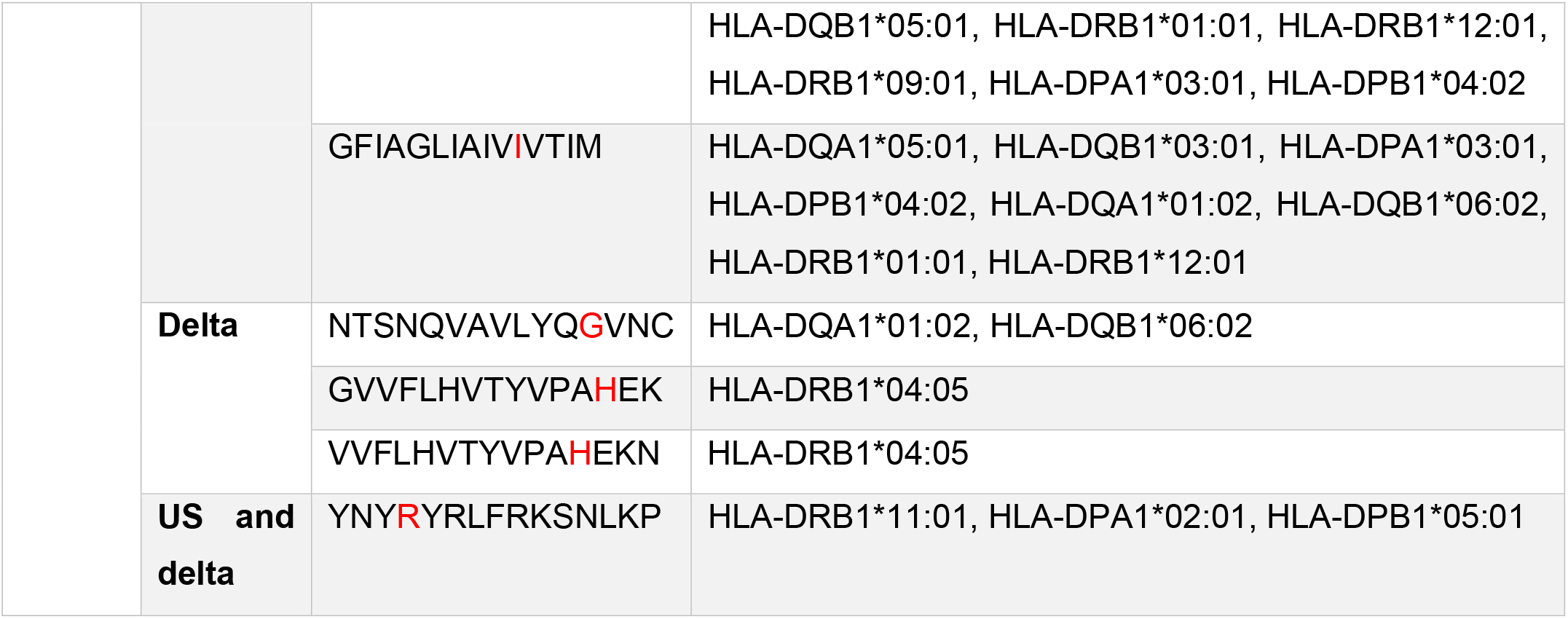
All antigenic non-toxic non-allergenic and stable CD4 epitopes per mutation for each SARS-CoV-2 variant N and S protein, with the variant-specific mutations written in red.

Despite the success of current vaccines, it remains necessary to expand the tools at our disposal to combat the current COVID-19 pandemic and prepare for future variants and outbreaks. Specifically, epitope-based vaccines present a useful tool with advantages over the current mRNA vaccines. Firstly, epitope-based vaccines may be able to confer immunity that lasts longer than natural immunity or immunity induced by current vaccines. Natural immunity from previous infection has been shown to be unreliable in older individuals [70] and fades relatively quickly with a half-life of 200 days for T cells [71], making it ineffective in providing long term immunity against reinfection, with T cell immunity fading faster than B cell immunity [72]. Epitope-based vaccines can be designed to target every aspect of the immune system by including epitopes that activate both CD8 and CD4 T cells as well as B cells, all of which are necessary to confer long-term immunity. While all current COVID-19 vaccines only include the spike protein of SARS-CoV-2, it has been shown that only N protein T cell epitopes for SARS-CoV lasted at least 17 years, making them the most long-lasting epitopes [32]. This suggests an opportunity to generate longer lasting immunity by making use of the longevity of N protein epitopes in vaccines. Multi-epitope vaccines can include epitopes of both the S and N proteins of SARS-CoV-2, such as those suggested by our study, to provide the most effective and long-lasting protection against SARS-CoV-2. More studies are also needed to investigate how to prolong T cell immunity in a potentially universal coronavirus vaccine.

Another advantage of epitope-based vaccines is their specificity, which provides them with a unique ability to precisely target certain areas of a protein of interest. This ability is especially important in the context of an evolving virus such as SARS-CoV-2 as novel, increasingly infectious variants are spreading and continue to emerge. Epitope-based vaccines can be designed to target certain variants by including epitopes containing variant-specific mutations, such as those identified by our study. This feature may be of particular use in booster shots designed to target specific variants against which the current vaccines may be less effective [73]. T cells, IgG and IgM antibodies have been shown to decrease substantially 6 months after administering the 2nd dose of the “low-dose” Moderna vaccine [74], suggesting that booster shots may be necessary to confer long-lasting immunity by reactivating cellular and humoral immunity. Of particular concern currently is the Delta variant that is spreading globally, as recent data suggests that both the AstraZeneca and Pfizer vaccines are slightly less effective against this variant in preventing COVID-19 related symptoms compared to the Alpha variant [75]. A recent large-scale study of Delta infections in the UK found that the protection offered against the variant by the AstraZeneca and Pfizer vaccines wanes with time, further supporting the need for variant-specific vaccines to boost immunity against variants of concern [76]. Despite global vaccination efforts, the rise in COVID-19 cases around the world as the Delta variant of SARS-CoV-2 spreads implies that variant-specific booster shots may be needed to combat emerging variants. Our study identified high-quality epitopes specific to current variants of concern that can be included in epitope-based vaccines to target most variants of concern.

The specificity of epitope-based vaccines also provides a potential method for preventing Original Antigenic Sin (OAS). According to this theory, the development of immunity against a pathogen is shaped by first exposure to that pathogen, limiting the flexibility of the immune system [77]. Upon subsequent infection with a novel strain of the pathogen, the immune system produces antibodies to the original strain it was first exposed to rather than mounting a response to novel antigens of the new strain. This phenomenon has been seen in response to influenza vaccines, demonstrating how OAS can allow novel influenza strains to evade the immune system despite vaccination [78]. OAS has also been seen in T cells, impairing the ability of cytotoxic T lymphocytes to respond effectively to variant viruses [79]. OAS is an important consideration in developing vaccines against SARS-CoV-2, as it suggests that immunity developed in response to infection or vaccination with one strain may not protect against all novel variants. Additionally, a booster vaccine that includes epitopes from the original reference sequence, such as using a variant of the SARS-CoV-2 spike protein, will likely reactivate B and T cells against the original epitopes without mounting a variant-specific immune response. For booster vaccines to induce a variant-specific immune response that can confer additional immunity against novel variants, they must be carefully formulated to avoid inducing an immune response against the original strain, a technique which requires great specificity. While some methods have been suggested to alleviate OAS [80], we hereby hypothesize a new method of preventing OAS through the use of epitope-based booster vaccines. We suggest that the specificity of epitope-based booster vaccines may allow them to prevent OAS and better activate variant-specific immune cells, thus providing stronger immunity against novel SARS-CoV-2 variants.

## Conclusion

In this study, we identified the best epitope prediction tools available on IEDB for predicting CD8 and CD4 binding epitopes of SARS-CoV-2. We then used these tools to predict SARS-CoV-2 N and S protein T cell epitopes. Prioritizing epitopes that were found to be common across the current SARS-CoV-2 variants, predicted to be immunogenic, non-toxic, non-allergenic, with high cell penetrability and with a high population coverage based on HLA alleles allowed us to identify high-quality T cell epitopes. We recommend these epitopes for a multi-epitope vaccine against the common SARS-CoV-2 variants, including variants of concern. We also present variant-specific immunogenic CD8 and CD4 T cell epitopes that can more precisely target variants of concern. Given the rise of increasingly infectious variants with the potential to evade natural immunity and current vaccines, these epitopes may prove useful in developing variant-specific booster shots to combat the most concerning variants. Together, these predicted epitopes may provide both broad protection against common SARS-CoV-2 variants, as well as more targeted protection against specific variants. While other studies have attempted to use immunoinformatics to design epitope-based vaccines against SARS-CoV-2, our comparison of available epitope prediction tools to experimentally validated SARS-CoV-2 epitopes validates that our predicted epitopes come from the most accurate available tool on IEDB.org. Additionally, our comparison of predicted epitopes across all current variants increases confidence that the effectiveness of our designed vaccine will not be undermined by the mutations of certain variants. Our epitope prediction together with our mouse MHC affinity prediction warrant further preclinical studies to validate their efficacy for use in designing epitope-based vaccines against SARS-CoV-2. While this experimental validation remains necessary, immunoinformatics has allowed us to predict epitopes in silico that are most likely to prove effective in vitro and in vivo. This will greatly reduce the cost and time of traditional vaccine development as it is urgent to develop new therapeutics and vaccines to combat this rapidly evolving pandemic as efficiently as possible.

## Supporting information

Supplementary Matarial 4

Supplementary Matarial 3

Supplementary Matarial 1

Supplementary Matarial 2

## Acknowledgements

We wish to acknowledge for the support by Lombardi Comprehensive Cancer Center, Georgetown University Medical Center.

