## Supplementary Matarial 1 for "Design of T cell epitope-based vaccine candidate for SARS-CoV-2 targeting nucleocapsid and spike protein escape variants"

**Supplementary Methods**

**Retrieval of N protein SARS-CoV-2 experimentally determined epitopes**

Finding CD8 experimental epitopes:

We obtained 51 epitopes and added 6 new epitopes: 7KGT, 7LG2, 7LG3, 7LFZ, 7KGR, 7KGS from RCSB.org. Then, we selected the 3 N protein epitopes that had at least 60% affinity to their MHC1 (Prachar et al), but they were all duplicates. We then obtained 1 from^7^ Sohail et al (6 in total, 5 duplicates), 1 from^8^ Schulien et al, 5 from^9^ Nelde et al, 5^10^ from Kared et al, 4^11^ from Saini et al and none^13^ from Tarke et al. Next, we used the immunome browser tool on IEDB.org and selected the N protein SARS-CoV-2 MHC1 epitopes that fit our criteria, have at least one positive assay and matched 100% with the reference N protein sequence. We obtained 15 new experimental epitopes. Next, we went to the ViPR.org website to obtain 76 N protein experimental epitopes for betacoronaviridae, most of which were for the old SARS virus and all 76 were already present in our repertoire of experimental epitopes. We ended up with 88 high-quality, experimentally determined SARS-CoV-2 N protein CD8 T cell epitopes.

**Choosing the best tool**

CD8 epitopes:

To assess the success of each tool, the predictions were matched with experimentally determined epitopes in 3 ways: lenient, intermediate and stringent. In the lenient method, if a predicted epitope is contained within an experimental epitope or vice versa, the two epitopes are matched. In the intermediate method, if the levenshtein score between the predicted and experimental epitopes is less than or equal to 1, the two epitopes match. In the stringent method, only matches of 100% are considered.

**Population coverage analysis**

Only HLA alleles from the top 300 predicted epitopes ordered according to “rank” were considered.

| Protein | Population Coverage | Epitopes | Common HLA Restriction | Mouse MHC Restriction | Instability Index |
| --- | --- | --- | --- | --- | --- |
| N | 98.46% | LSPRWYFYY | HLA-A*01:01, HLA-A*30:02, HLA-B*57:01 | H2-Dd | unstable |
|  |  | SPRWYFYYL | HLA-B*07:02, HLA-B*08:01 | Not available | unstable |
|  |  | DLSPRWYFY | HLA-A*26:01, HLA-A*01:01, HLA-A*30:02, HLA-A*33:01 | Not available | unstable |
|  |  | TPSGTWLTY | HLA-B*35:01, HLA-B*53:01, HLA-A*26:01, HLA-A*01:01 | Not available | stable |
|  |  | NTASWFTAL | HLA-A*68:02, HLA-A*26:01, HLA-A*02:06 | H2-Dd | stable |
|  |  | YYRRATRRI | HLA-A*24:02, HLA-A*23:01 | Not available | unstable |
|  |  | GYYRRATRR | HLA-A*31:01, HLA-A*33:01 | Not available | unstable |
|  |  | TGPEAGLPY | HLA-B*35:01 | H2-Dd | unstable |
|  |  | NNAAIVLQL | HLA-A*68:02 | Not available | stable |
|  |  | KTFPPTEPK | HLA-A*11:01, HLA-A*03:01, HLA-A*30:01, HLA-A*31:01, HLA-A*68:01, HLA-A*32:01 | Not available | unstable |
|  |  | NVTQAFGRR | HLA-A*68:01, HLA-A*33:01 | Not available | unstable |
|  |  | NFGDQELIR | HLA-A*33:01 | Not available | stable |
|  |  | GMSRIGMEV | HLA-A*02:03, HLA-A*02:01 | Not available | unstable |
|  |  | ELIRQGTDY | HLA-A*26:01, HLA-B*15:01 | Not available | stable |
|  |  | LPNNTASWF | HLA-B*53:01, HLA-B*35:01, HLA-B*07:02 | Not available | stable |
|  |  | KDLSPRWYF | HLA-B*44:02, HLA-A*32:01, HLA-B*44:03 | Not available | unstable |
|  |  | AGLPYGANK | HLA-A*30:01, HLA-A*11:01 | Not available | stable |
|  |  | EVTPSGTWL | HLA-A*68:02, HLA-A*26:01 | Not available | stable |
|  |  | ATKAYNVTQ | HLA-A*30:01 | Not available | stable |
|  |  | DAALALLLL | HLA-B*51:01, HLA-A*68:02 | Not available | stable |
|  |  | RIRGGDGKM | HLA-B*07:02 | Not available | stable |
|  |  | AYNVTQAFGR | HLA-A*31:01 | Not available | stable |
|  |  | DLSPRWYFYY | HLA-A*26:01 | Not available | unstable |
|  |  | EAGLPYGANK | HLA-A*68:01 | Not available | stable |
|  |  | KDLSPRWYFY | HLA-A*30:02 | Not available | unstable |
|  |  | KTFPPTEPKK | HLA-A*03:01, HLA-A*11:01, HLA-A*30:01, HLA-A*31:01, HLA-A*68:01 | Not available | unstable |
|  |  | MEVTPSGTWL | HLA-B*40:01, HLA-B*44:03, HLA-B*44:02 | Not available | stable |
|  |  | QELIRQGTDY | HLA-B*44:03, HLA-B*44:02 | Not available | stable |
|  |  | SASAFFGMSR | HLA-A*68:01 | Not available | unstable |
|  |  | VTPSGTWLTY | HLA-A*01:01, HLA-B*35:01, HLA-A*30:02, HLA-B*53:01, HLA-A*26:01 | Not available | stable |
|  |  | YKTFPPTEPK | HLA-A*11:01, HLA-A*68:01, HLA-A*03:01, HLA-A*30:01 | Not available | unstable |
| S | 92.02% | ADAGFIKQY | HLA-B*44:02, HLA-B*44:03 | Not available | stable |
|  |  | AEIRASANL | HLA-B*40:01, HLA-B*44:03, HLA-B*44:02 | H2-Dd | stable |
|  |  | AEVQIDRLI | HLA-B*44:03, HLA-B*44:02, HLA-B*40:01 | Not available | stable |
|  |  | DAVRDPQTL | HLA-B*51:01 | H2-Db | stable |
|  |  | FIAGLIAIV | HLA-A*02:03, HLA-A*02:06 | Not available | stable |
|  |  | FTISVTTEI | HLA-A*68:02, HLA-A*02:06 | H2-Db | unstable |
|  |  | GEVFNATRF | HLA-B*44:03, HLA-B*44:02 | H2-Db | stable |
|  |  | HWFVTQRNF | HLA-A*23:01, HLA-A*24:02 | Not available | stable |
|  |  | INITRFQTL | HLA-B*08:01 | H2-Db | unstable |
|  |  | KEIDRLNEV | HLA-B*40:01 | Not available | stable |
|  |  | LAGTITSGW | HLA-B*58:01, HLA-B*57:01, HLA-B*53:01 | Not available | stable |
|  |  | LPFNDGVYF | HLA-B*35:01, HLA-B*53:01, HLA-B*51:01 | Not available | unstable |
|  |  | LTDEMIAQY | HLA-A*01:01, HLA-A*30:02 | Not available | stable |
|  |  | NASVVNIQK | HLA-A*68:01 | Not available | unstable |
|  |  | NTQEVFAQV | HLA-A*68:02 | Not available | stable |
|  |  | PYRVVVLSF | HLA-A*23:01, HLA-A*24:02 | Not available | stable |
|  |  | QLTPTWRVY | HLA-A*30:02, HLA-B*15:01 | Not available | stable |
|  |  | QPRTFLLKY | HLA-B*35:01 | Not available | stable |
|  |  | RSFIEDLLF | HLA-B*58:01, HLA-B*57:01, HLA-A*32:01 | Not available | unstable |
|  |  | SANNCTFEY | HLA-A*30:02, HLA-B*35:01 | Not available | unstable |
|  |  | VVFLHVTYV | HLA-A*02:03, HLA-A*02:06, HLA-A*68:02, HLA-A*02:01 | H2-Db | stable |
|  |  | WTAGAAAYY | HLA-A*26:01, HLA-A*01:01 | Not available | stable |
|  |  | YQPYRVVVL | HLA-B*08:01 | H2-Dd | stable |
|  |  | EILDITPCSF | HLA-A*26:01 | Not available | stable |
|  |  | KLNDLCFTNV | HLA-A*02:03 | Not available | stable |
|  |  | KSFTVEKGIY | HLA-A*30:02 | Not available | stable |
|  |  | LADAGFIKQY | HLA-A*01:01 | Not available | stable |
|  |  | LPIGINITRF | HLA-B*53:01, HLA-B*35:01 | H2-Ld | unstable |
|  |  | NFTISVTTEI | HLA-A*68:02 | H2-Kd | stable |
|  |  | SSANNCTFEY | HLA-A*01:01 | Not available | unstable |
|  |  | VLPFNDGVYF | HLA-B*53:01 | H2-Db | stable |

**Table S1.** All CD8 common N and S protein non-toxic non-allergenic epitopes across all studied SARS-CoV-2 variants and their world population coverage.

| Protein | Population Coverage | Epitopes | Common HLA restriction | Mouse MHC |
| --- | --- | --- | --- | --- |
| N | 78.28% | TPSGTWLTY | HLA-B*35:01, HLA-B*53:01, HLA-A*26:01, HLA-A*01:01 | H-2-Ld |
|  |  | NTASWFTAL | HLA-A*68:02, HLA-A*26:01, HLA-A*02:06 | H-2-Kb, H-2-Dd |
|  |  | NNAAIVLQL | HLA-A*68:02 | Not available |
|  |  | NFGDQELIR | HLA-A*33:01 | Not available |
|  |  | ELIRQGTDY | HLA-A*26:01, HLA-B*15:01 | Not available |
|  |  | LPNNTASWF | HLA-B*53:01, HLA-B*35:01, HLA-B*07:02 | H-2-Ld |
|  |  | AGLPYGANK | HLA-A*30:01, HLA-A*11:01 | Not available |
|  |  | EVTPSGTWL | HLA-A*68:02, HLA-A*26:01 | Not available |
|  |  | ATKAYNVTQ | HLA-A*30:01 | Not available |
|  |  | DAALALLLL | HLA-B*51:01, HLA-A*68:02 | Not available |
|  |  | RIRGGDGKM | HLA-B*07:02 | Not available |
|  |  | AYNVTQAFGR | HLA-A*31:01 | Not available |
|  |  | EAGLPYGANK | HLA-A*68:01 | Not available |
|  |  | MEVTPSGTWL | HLA-B*40:01, HLA-B*44:03, HLA-B*44:02 | Not available |
|  |  | VTPSGTWLTY | HLA-A*01:01, HLA-B*35:01, HLA-A*30:02, HLA-B*53:01, HLA-A*26:01 | Not available |
| S | 87.13% | ADAGFIKQY | HLA-B*44:02, HLA-B*44:03 | Not available |
|  |  | AEIRASANL | HLA-B*40:01, HLA-B*44:03, HLA-B*44:02 | H-2-Dd |
|  |  | AEVQIDRLI | HLA-B*44:03, HLA-B*44:02, HLA-B*40:01 | Not available |
|  |  | DAVRDPQTL | HLA-B*51:01 | H-2-Db, H-2-Dd, H-2-Kd |
|  |  | FIAGLIAIV | HLA-A*02:03, HLA-A*02:06 | Not available |
|  |  | GEVFNATRF | HLA-B*44:03, HLA-B*44:02 | H-2-Db |
|  |  | HWFVTQRNF | HLA-A*23:01, HLA-A*24:02 | Not available |
|  |  | KEIDRLNEV | HLA-B*40:01 | Not available |
|  |  | LAGTITSGW | HLA-B*58:01, HLA-B*57:01, HLA-B*53:01 | H-2-Ld |
|  |  | LTDEMIAQY | HLA-A*01:01, HLA-A*30:02 | Not available |
|  |  | NTQEVFAQV | HLA-A*68:02 | Not available |
|  |  | PYRVVVLSF | HLA-A*23:01, HLA-A*24:02 | Not available |
|  |  | VVFLHVTYV | HLA-A*02:03, HLA-A*02:06, HLA-A*68:02, HLA-A*02:01 | H-2-Db, H-2-Kb, H-2-Dd |
|  |  | WTAGAAAYY | HLA-A*26:01, HLA-A*01:01 | Not available |
|  |  | YQPYRVVVL | HLA-B*08:01 | H-2-Dd |
|  |  | EILDITPCSF | HLA-A*26:01 | Not available |
|  |  | KLNDLCFTNV | HLA-A*02:03 | Not available |
|  |  | KSFTVEKGIY | HLA-A*30:02 | Not available |
|  |  | LADAGFIKQY | HLA-A*01:01 | Not available |
|  |  | NFTISVTTEI | HLA-A*68:02 | H2-Kd |
|  |  | VLPFNDGVYF | HLA-B*53:01 | H2-Db |

**Table S2.** CD8 common immunogenic, non-toxic, non-allergenic and stable N and S protein epitopes across all studied SARS-CoV-2 variants and their world population coverage.

| Protein | Variant | Epitope | HLA Restriction | Mouse MHC Restriction | Stability |
| --- | --- | --- | --- | --- | --- |
| N | **Beta** | GSSRGISPAR | HLA-A*31:01 | Not available | unstable |
|  | **Gamma** | SSRDDQIGYY | HLA-A*01:01, HLA-A*30:02, HLA-A*26:01 | Not available | unstable |
|  |  | SSRDDQIGY | HLA-A*30:02, HLA-A*01:01, HLA-B*15:01, HLA-A*26:01, HLA-B*35:01, HLA-A*30:01, HLA-B*57:01 | Not available | unstable |
|  | **Alpha** | SLNGPQNQR | HLA-A*31:01, HLA-A*03:01, HLA-A*33:01, HLA-A*68:01, HLA-A*11:01 | Not available | stable |
|  |  | MSLNGPQNQR | HLA-A*68:01, HLA-A*31:01, HLA-A*33:01 | Not available | unstable |
|  | **US-AUS** | RTSPARMAG | HLA-A*30:01 | Not available | unstable |
|  |  | RNSTPGSSK | HLA-A*30:01 | Not available | unstable |
|  |  | NSTPGSSKR | HLA-A*68:01, HLA-A*33:01 | Not available | unstable |
| S | **Beta** | TPINLVRGL | HLA-B*07:02 | H-2-Ld | stable |
|  |  | RFANPVLPF | HLA-A*24:02, HLA-A*23:01 | Not available | unstable |
|  | **Gamma** | GQTGTIADY | HLA-B*15:01 | Not available | stable |
|  |  | ASANLAAIK | HLA-A*11:01 | Not available | stable |
|  |  | YPFLGVYYH | HLA-B*35:01 | Not available | unstable |
|  |  | GTIADYNYK | HLA-A*11:01 | Not available | stable |
|  | **Alpha** | VLNDILARL | HLA-A*02:03, HLA-A*02:01, HLA-A*02:06 | H-2-Kb, H-2-Dd | stable |
|  |  | SVLNDILARL | HLA-A*02:03 | H-2-Kb | stable |
|  |  | QIITTHNTF | HLA-B*15:01 | H-2-Db, H-2-Dd, H-2-Kd, H-2-Ld | stable |
|  |  | SVLNDILAR | HLA-A*11:01 | Not available | stable |
|  | **US** | YNYRYRLFR | HLA-A*33:01 | Not available | stable |
|  |  | LPLVSIQCV | HLA-B*51:01 | H-2-Ld | stable |
|  | **Cluster 5** | YNYLFRLFR | HLA-A*33:01 | Not available | stable |
|  |  | NYNYLFRLFR | HLA-A*33:01 | Not available | stable |
|  | **Delta** | GVYFASIEK | HLA-A*11:01, HLA-A*03:01 | Not available | unstable |
|  |  | IEKSNIIRGW | HLA-B*44:02, HLA-B*44:03 | Not available | unstable |
|  |  | WMKSEFRVY | HLA-B*15:01 | Not available | unstable |
|  |  | KSWMKSEFR | HLA-A*31:01 | Not available | unstable |
|  |  | TYVPAHEKNF | HLA-A*24:02, HLA-A*23:01 | H-2-Kd | stable |
|  | **US and delta** | KVGGNYNYR | HLA-A*31:01 | Not available | stable |

**Table S3.** All variant-specific immunogenic, non-toxic and non-allergenic CD8 SARS-CoV-2 N and S protein epitopes with the variant-specific mutations written in red.

| Protein | Population Coverage | Epitopes | Common HLA restriction | Stability |
| --- | --- | --- | --- | --- |
| N | 45.40% | AQFAPSASAFFGMSR | HLA-DRB1*09:01 | unstable |
|  |  | GTRNPANNAAIVLQL | HLA-DQA1*01:02, HLA-DQB1*06:02 | stable |
|  |  | QIGYYRRATRRIRGG | HLA-DRB1*11:01 | unstable |
|  |  | GYYRRATRRIRGGDG | HLA-DRB1*11:01 | unstable |
|  |  | IGYYRRATRRIRGGD | HLA-DRB1*11:01 | unstable |
| S | 97.11% | YRVVVLSFELLHAPA | HLA-DPA1*03:01, HLA-DPB1*04:02, HLA-DPA1*02:01, HLA-DPB1*01:01, HLA-DPA1*01:03, HLA-DPB1*02:01 | stable |
|  |  | VVVLSFELLHAPATV | HLA-DRB1*01:01 | stable |
|  |  | VVLSFELLHAPATVC | HLA-DRB1*01:01 | stable |
|  |  | VLSFELLHAPATVCG | HLA-DRB1*01:01 | stable |
|  |  | TFEYVSQPFLMDLEG | HLA-DPA1*01:03, HLA-DPB1*04:01 | unstable |
|  |  | RVVVLSFELLHAPAT | HLA-DRB1*01:01 | stable |
|  |  | QSLLIVNNATNVVIK | HLA-DRB1*13:02, HLA-DRB3*02:02 | stable |
|  |  | NCTFEYVSQPFLMDL | HLA-DPA1*01:03, HLA-DPB1*04:01 | unstable |
|  |  | INITRFQTLLALHRS | HLA-DRB5*01:01 | unstable |
|  |  | GWTFGAGAALQIPFA | HLA-DRB1*09:01 | stable |
|  |  | CTFEYVSQPFLMDLE | HLA-DPA1*01:03, HLA-DPB1*04:01 | unstable |

**Table S4.** All CD4 common N and S protein antigenic non-toxic non-allergenic epitopes across all studied SARS-CoV-2 variants and their world population coverage.

| Protein | Variant | Epitope | HLA restriction | stability |
| --- | --- | --- | --- | --- |
| N | **Gamma** | RDDQIGYYRRATRRI | HLA-DRB3*01:01 | unstable |
|  |  | SSRDDQIGYYRRATR | HLA-DRB1*03:01 | unstable |
|  |  | SRDDQIGYYRRATRR | HLA-DRB1*03:01 | unstable |
|  | **US-AUS** | STPGSSKRTSPARMA | HLA-DRB1*09:01 | unstable |
|  | **Delta** | KKKKAYETQALPQRQ | HLA-DPA1*02:01, HLA-DPB1*14:01 | stable |
|  |  | DKKKKAYETQALPQR | HLA-DPA1*02:01, HLA-DPB1*01:01 | stable |
|  |  | KKKAYETQALPQRQK | HLA-DPA1*02:01, HLA-DPB1*01:01 | stable |
| S | **Beta** | HTPINLVRGLPQGFS | HLA-DRB4*01:01, HLA-DRB1*11:01, HLA-DRB1*08:02, HLA-DRB1*04:05 | stable |
|  | **Gamma** | LVLLPLVSSQCVNFT | HLA-DRB4*01:01, HLA-DRB1*08:02, HLA-DRB1*04:01, HLA-DRB1*15:01, HLA-DRB1*12:01, HLA-DRB1*04:05 | stable |
|  |  | FLVLLPLVSSQCVNF | HLA-DRB1*01:01, HLA-DRB4*01:01, HLA-DRB1*08:02, HLA-DRB1*11:01, HLA-DRB1*12:01, HLA-DRB1*04:05, HLA-DRB1*15:01, HLA-DRB5*01:01, HLA-DRB1*04:01 | stable |
|  |  | VLLPLVSSQCVNFTN | HLA-DRB4*01:01, HLA-DRB1*04:05 | stable |
|  |  | LLPLVSSQCVNFTNR | HLA-DRB4*01:01 | stable |
|  |  | LPLVSSQCVNFTNRT | HLA-DRB4*01:01 | stable |
|  |  | AEYVNNSYECDIPIG | HLA-DQA1*01:01, HLA-DQB1*05:01 | unstable |
|  |  | EYVNNSYECDIPIGA | HLA-DQA1*01:01, HLA-DQB1*05:01, HLA-DRB3*01:01 | stable |
|  |  | GAEYVNNSYECDIPI | HLA-DQA1*01:01, HLA-DQB1*05:01 | stable |
|  | **Alpha** | KKFLPFQQFGRDIDD | HLA-DPA1*02:01, HLA-DPB1*01:01, HLA-DPA1*02:01, HLA-DPB1*05:01, HLA-DRB5*01:01, HLA-DPA1*01:03, HLA-DPB1*04:01, HLA-DPA1*01:03, HLA-DPB1*02:01 | unstable |
|  |  | HRRARSVASQSIIAY | HLA-DPA1*02:01, HLA-DPB1*14:01, HLA-DRB1*07:01, HLA-DRB4*01:01 | unstable |
|  |  | AIPINFTISVTTEIL | HLA-DRB1*07:01, HLA-DRB1*09:01, HLA-DRB3*02:02, HLA-DQA1*03:01, HLA-DQB1*03:02, HLA-DRB1*04:01, HLA-DRB1*13:02, HLA-DQA1*04:01, HLA-DQB1*04:02, HLA-DQA1*05:01, HLA-DQB1*02:01, HLA-DRB1*04:05, HLA-DRB3*01:01 | stable |
|  |  | INFTISVTTEILPVS | HLA-DRB1*07:01, HLA-DRB1*09:01, HLA-DRB3*02:02, HLA-DQA1*03:01, HLA-DQB1*03:02, HLA-DQA1*05:01, HLA-DQB1*02:01, HLA-DPA1*02:01, HLA-DPB1*01:01, HLA-DPA1*02:01, HLA-DPB1*14:01, HLA-DRB1*04:01, HLA-DPA1*03:01, HLA-DPB1*04:02, HLA-DPA1*01:03, HLA-DPB1*04:01, HLA-DRB3*01:01 | unstable |
|  | **US** | FVFLVLLPLVSIQCV | HLA-DRB1*01:01, HLA-DPA1*03:01, HLA-DPB1*04:02, HLA-DRB1*11:01, HLA-DPA1*02:01, HLA-DPB1*01:01, HLA-DRB5*01:01, HLA-DRB1*04:01, HLA-DRB1*15:01 | stable |
|  |  | MFVFLVLLPLVSIQC | HLA-DRB1*01:01, HLA-DPA1*03:01, HLA-DPB1*04:02, HLA-DRB1*11:01, HLA-DPA1*02:01, HLA-DPB1*01:01, HLA-DRB5*01:01, HLA-DRB1*04:01, HLA-DRB1*15:01, HLA-DPA1*01:03, HLA-DPB1*02:01 | stable |
|  |  | VLLPLVSIQCVNLTT | HLA-DRB4*01:01 | stable |
|  |  | LVLLPLVSIQCVNLT | HLA-DRB4*01:01 | stable |
|  |  | LGVYYHKNNKSCMES | HLA-DRB3*02:02 | unstable |
|  | **Cluster 5** | IWLGFIAGLIAIVIV | HLA-DQA1*05:01, HLA-DQB1*03:01, HLA-DRB1*01:01, HLA-DRB1*12:01, HLA-DQA1*01:01, HLA-DQB1*05:01, HLA-DRB1*09:01, HLA-DPA1*03:01, HLA-DPB1*04:02 | stable |
|  |  | YIWLGFIAGLIAIVI | HLA-DQA1*05:01, HLA-DQB1*03:01, HLA-DQA1*01:01, HLA-DQB1*05:01, HLA-DRB1*01:01, HLA-DRB1*12:01, HLA-DRB1*09:01, HLA-DPA1*03:01, HLA-DPB1*04:02 | stable |
|  |  | GFIAGLIAIVIVTIM | HLA-DQA1*05:01, HLA-DQB1*03:01, HLA-DPA1*03:01, HLA-DPB1*04:02, HLA-DQA1*01:02, HLA-DQB1*06:02, HLA-DRB1*01:01, HLA-DRB1*12:01 | stable |
|  | **Delta** | NGVQGFNCYFPLQSY | HLA-DQA1*01:01, HLA-DQB1*05:01 | unstable |
|  |  | NTSNQVAVLYQGVNC | HLA-DQA1*01:02, HLA-DQB1*06:02 | stable |
|  |  | RRRARSVASQSIIAY | HLA-DPA1*02:01, HLA-DPB1*14:01, HLA-DRB1*07:01 | unstable |
|  |  | SRRRARSVASQSIIA | HLA-DPA1*02:01, HLA-DPB1*14:01, HLA-DRB1*07:01 | unstable |
|  |  | NSRRRARSVASQSII | HLA-DPA1*02:01, HLA-DPB1*14:01, HLA-DRB1*07:01 | unstable |
|  |  | GVVFLHVTYVPAHEK | HLA-DRB1*04:05 | stable |
|  |  | VVFLHVTYVPAHEKN | HLA-DRB1*04:05 | stable |
|  | **US and Delta** | YNYRYRLFRKSNLKP | HLA-DRB1*11:01, HLA-DPA1*02:01, HLA-DPB1*05:01 | stable |

**Table S5.**  All variant-specific antigenic non-toxic non-allergenic CD4 SARS-CoV-2 N and S protein epitopes with the variant-specific mutations written in red.

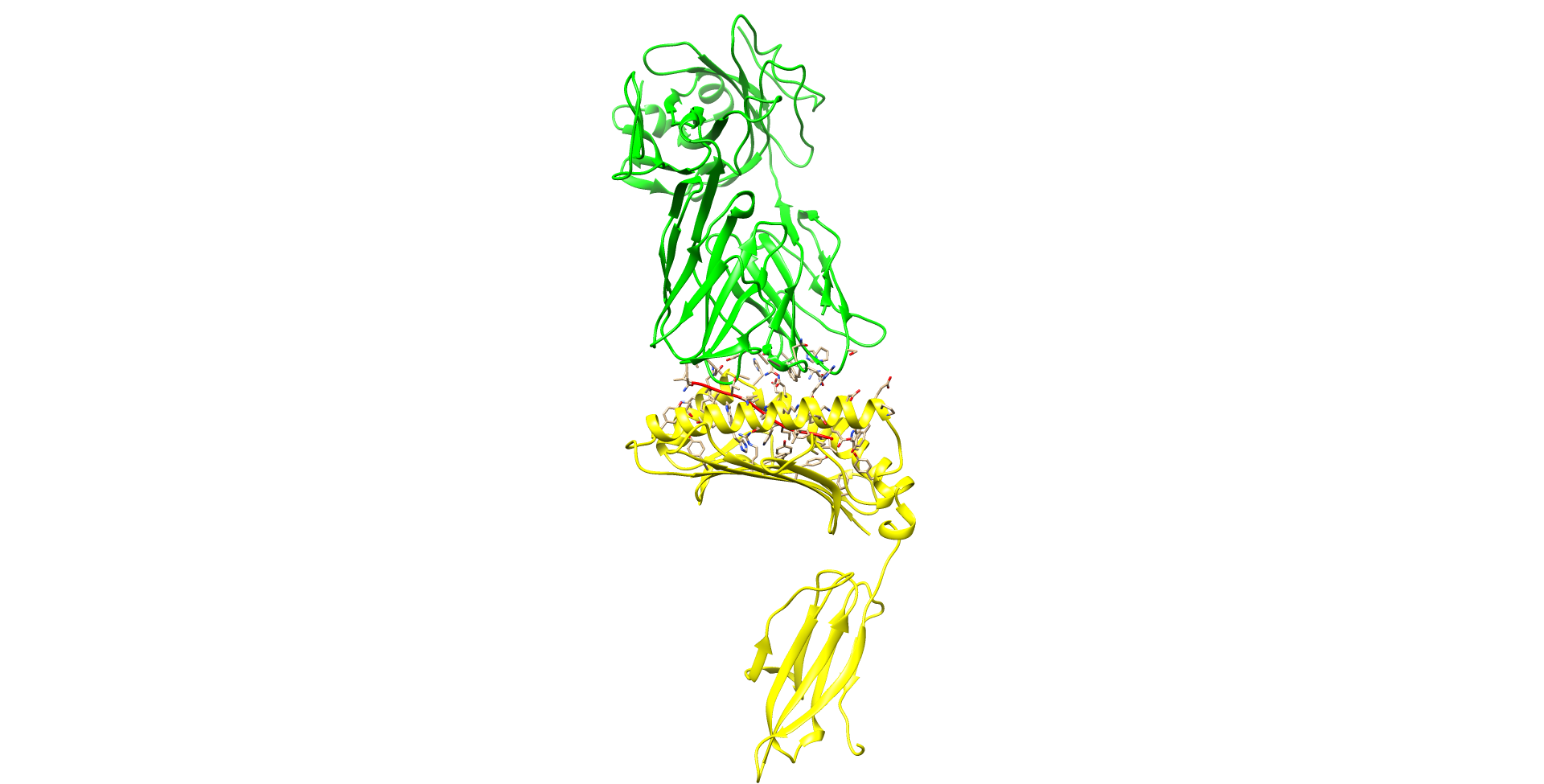

**Figure S1.** Long shot of the predicted 3D structure for the interaction of the CD8 S protein epitope VVFLHVTYV with HLA-A*02:01 and pRLQ3 TCR.
